# Mouse models of *NADK2* deficiency analyzed for metabolic and gene expression changes to elucidate pathophysiology

**DOI:** 10.1101/2021.12.10.472125

**Authors:** G. Murray, P. Bais, C. Hatton, A.L.D. Tadenev, K.H. Morelli, D. Schroeder, R. Doty, O. Fiehn, S.W.M. John, C.J. Bult, G.A. Cox, R.W. Burgess

## Abstract

*NADK2* encodes the mitochondrial isoform of NAD Kinase, which phosphorylates nicotinamide adenine dinucleotide (NAD). Rare recessive mutations in human *NADK2* are associated with a syndromic neurological mitochondrial disease that includes metabolic changes such as hyperlysinemia and 2,4 dienoyl CoA reductase (DECR) deficiency. However, the full pathophysiology resulting from *NADK2* deficiency is not known. Here we describe two chemically-induced mouse mutations in *Nadk2*, S326L and S330P, which cause a severe neuromuscular disease and shorten lifespan. The S330P allele was characterized in detail and shown to have marked denervation of neuromuscular junctions by 5 weeks of age and muscle atrophy by 11 weeks of age. Cerebellar Purkinje cells also showed progressive degeneration in this model. Transcriptome profiling on brain and muscle was performed at early and late disease stages. In addition, metabolomic profiling was performed on brain, muscle, liver, and spinal cord at the same ages. Combined transcriptomic and metabolomic analyses identified hyperlysinemia, DECR deficiency, and generalized metabolic dysfunction in *Nadk2* mutant mice, indicating relevance to the human disease. We compared findings from the *Nadk* model to equivalent RNAseq and metabolomic datasets from a mouse model of infantile neuroaxonal dystrophy, caused by recessive mutations in *Pla2g6*. This enabled us to identify disrupted biological processes that are common between these mouse models of neurological disease, such as translation, and those processes that are gene-specific such as glycolysis and acetylcholine binding. These findings improve our understanding of the pathophysiology of both *Nadk2* and *Pla2g6* mutations, as well as pathways common to neuromuscular/neurodegenerative diseases.

## Introduction

The pyridine nucleotide nicotinamide adenine dinucleotide (NAD) and its phosphorylated form, NADP, are found in all organisms and are involved in a myriad of biological functions, including serving as substrates and cofactors for numerous enzymes and metabolic processes. NAD is converted to NADP by NAD kinases (NADKs). Since NAD and NADP are membrane impermeant, these kinases are compartmentalized within the cell (1). The cytosolic form of human NADK was identified in 2001 (2), but it was not until 2012 that the mitochondrial form of NADK was discovered (NADK2, also called MNADK for mitochondrial NAD kinase, and *C5ORF33*, prior to the functional characterization of the human gene product) (3). The initial characterization of NADK2 showed that this kinase phosphorylates NAD using ATP or inorganic polyphosphate as a substrate, and localizes to mitochondria due to an amino-terminal mitochondrial targeting sequence (3).

In human genetic studies, three patients have been described with recessive *NADK2* mutations, as summarized below. The first had hyperlysinemia and 2,4 dienoyl CoA reductase (DECR) deficiency, indicated by elevated levels of C10:2 carnitine (4). At eight-weeks-of-age, the patient had failure to thrive, microcephaly, central hypotonia and dysmorphic features. With age, neurological features increased with encephalopathy, dystonia, spastic quadriplegia, and epilepsy. The patient died at 5 years-of-age. Whole exome sequencing revealed a homozygous R340X truncating mutation in *NADK2* in this patient. One previous patient with similar presentation, but even earlier mortality, was described in 1990, but no genetic results were reported (5).

In the second patient described, the general metabolic and neurological presentation were very similar to those described above (6). However, the patient was placed on a lysine-restricted diet at one-month-of-age, and treatment with pyridoxal phosphate at three-years-of-age improved electroclinical symptoms. At age ten, the patient had ataxia, poor coordination, cerebellar atrophy, and oromotordysphasia, but understood fluid speech and was social. She was diagnosed as having a similar homozygous truncating mutation, W319Cfs*21.

The third patient has much milder symptoms and later onset (7). She was ascertained at nine-years-of-age with normal intelligence, but decreased visual acuity, optic atrophy, nystagmus, and peripheral neuropathy. Hyperlysinemia was present, as well as hyperprolinemia, but C10:2 carnitine levels were normal. Sequencing through the Undiagnosed Disease Network identified a homozygous Met1Val change in *NADK2. NADK2* has multiple transcripts, though initiation of translation at the next downstream ATG is not expected to produce a protein with a mitochondrial targeting sequence. Consistent with this, patient fibroblasts revealed residual protein expression, though primarily in the cytosolic fraction. This, and the milder phenotype, suggest this is a hypomorphic allele of *NADK2*, producing protein through an unknown mechanism.

Hyperlysinemia is caused by mutations in *AASS*, encoding α-aminoadipic semialdehyde synthase, which degrades lysine (8,9). Similarly, defects in DECR cause elevated C10:2 carnitine due to changes in beta-oxidation of polyunsaturated fatty acids (10). Both AASS and DECR enzymatic activities depend on NADP as a co-substrate. Furthermore, in the absence of mitochondrial NADP due to *NADK2* mutations, these enzymes are either inactive (DECR1, the mitochondrial DECR enzyme, DECR2 localizes to peroxisomes), or reduced in their levels (AASS), suggesting a molecular chaperone role for NADP (4). Thus, the molecular basis for hyperlysinemia and C10:2 carnitine elevation in NADK2-deficient patients may be understood. However, AASS deficiency is considered a benign metabolic variant (8,11), and mice lacking *Decr1* are viable and have sensitivity to fasting and cold stress, but no reported neurological phenotype or shortened lifespan (12,13). Therefore, these pathways alone are unlikely to account for the severe disease observed in patients with mutations in *NADK2*, and a deeper analysis of the pathophysiology of this rare disorder is needed.

Model organisms have the potential to help elucidate the pathophysiology of *NADK2* deficiency. Two independent null alleles of mouse *Nadk2, tm1b (EUCOMM)Wtsi* (MGI:5637034) and *em1(IMPC)J* (MGI:5689888), produced and analyzed through the KOMP (Knockout Mouse Program), are early lethal with complete penetrance. The *em1J* homozygotes die at 12.5 days of gestation (E12.5), indicating null alleles of *Nadk2* are embryonic lethal in mice. Analysis has also been performed on the *tm1a(EUCOMM)Wtsi* allele (MGI:4842252), which includes a *LacZ* reporter and gene-trapping insertion, but does not include the deletion of a critical, frame shifting exon as in the *tm1b* allele, and may therefore result in a hypomorph (https://www.infrafrontier.eu/sites/infrafrontier.eu/files/upload/public/pdf/Resources%20and%20Services/eucomm-alleles-overview_infrafrontier-2016.pdf). The homozygous *Nadk2^tm1a(EUCOMM)Wtsi^* mice survive and have relevant elevated lysine and C10:2 carnitine levels (14). They were analyzed primarily for liver metabolic phenotypes and develop non-alcoholic fatty liver when fed a high-fat diet. No neurological symptoms or shortened lifespan were reported.

Here we describe two chemically-induced point mutations in *Nadk2* that present with clear neuromuscular phenotypes, neurodegeneration, shortened lifespan, and metabolic deficits. To better understand the pathophysiology underlying these phenotypes, we performed extensive metabolomic and transcriptomic analyses in these mice, and compared them to mice carrying a mutation in *Pla2g6* (15), a model of infantile neuroaxonal dystrophy (INAD, (16)), to separate changes associated with general neuromuscular dysfunction from gene- and disease-specific effects.

## Results

### Identification of *Nadk2* mouse mutations

We identified two independent strains of mice with mutations in *Nadk2*. The mice were first analyzed because of their overt neuromuscular phenotypes, including hind limb wasting, gait abnormalities, and shakiness. In addition, while a formal lifespan study was not performed, affected mice begin dying at 3-4 months of age, indicating early mortality. The mutations were produced in chemical mutagenesis programs performed at The Jackson Laboratory and were recessive (unaffected parents produced ~25% affected offspring). They were mapped using standard positional cloning strategies (see methods), and both were located on proximal Chr. 15 (Figure 1A). The first mutation was identified in the *Nadk2* gene, located at 9.1 mb on Chr. 15 (designated *Nadk2^m1Jcs^*, MGI:6825802) by sequencing PCR amplified cDNA of candidate genes in the interval, which revealed a homozygous *Nadk2* mutation, changing Serine 326 to Leucine (S326L) (Figure 1B). Given the overlapping map positions and similar phenotypes of the two mutations, we sequenced the *Nadk2* gene in the second strain (*Nadk2^nmf421^*, MGI:6825801), and also identified a homozygous *Nadk2* mutation changing Serine 330 to Proline (S330P) (Figure 1B). To confirm these mutations were causal, we performed a complementation test, breeding mice heterozygous for each mutation to produce compound heterozygotes (S326L/S330P). This produced affected offspring with a similar phenotype, including two mice that were small and shaky and died by ~3 months of age and a third with a similar overt phenotype. These mice were confirmed to be compound heterozygotes by genotyping. Thus, we have identified two independent mutations in the *Nadk2* gene in mice that cause an overt phenotype and shortened lifespan. Given the similar phenotype of the two strains, only the *nmf421* S330P allele was used in the experiments described below and homozygotes are referred to as *Nadk2^-/-^* for simplicity. The *m1Jcs* S326L allele is cryopreserved.

**Figure 1.**
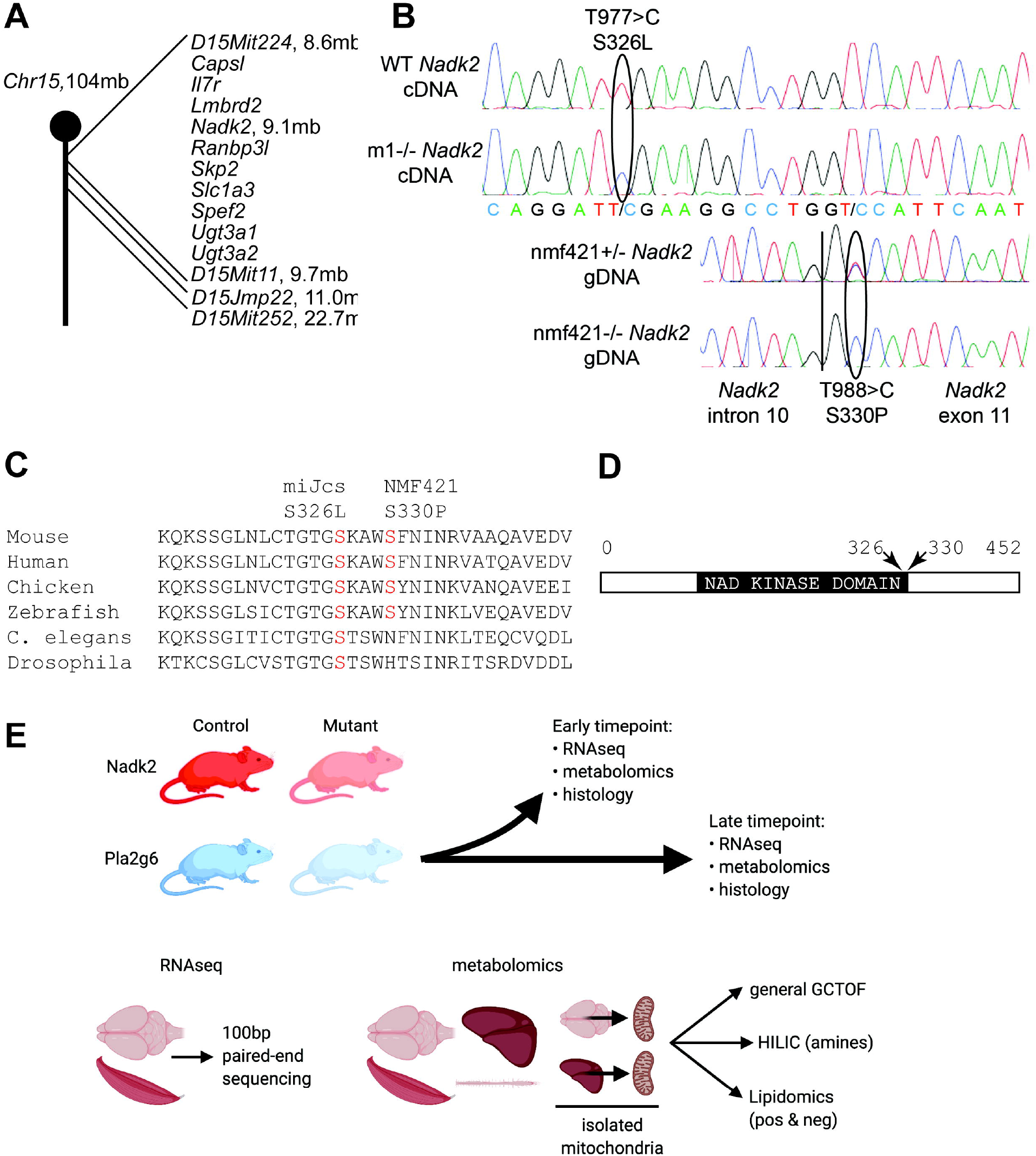
*Nadk2* mouse mutations and study design. A) The ENU-induced mutations mapped to proximal mouse Chr. 15, with the *m1Jcs* allele mapping between D15Mit224 at 8.6 mb and D15Mit11 at 9.7 mb. This interval contains ten protein coding genes. The *nmf421* allele mapped proximally to D15Jmp22 at 11.0 mb. B) Sequencing chromatograms showing the T977>C mutation in cDNA prepared from wild type and *m1Jcs* homozygous mice, and the T988>C mutation in PCR-amplified genomic DNA from heterozygous and homozygous *nmf421* mice. The vertical line in genomic DNA sequence is the intron 10/exon 11 boundary. C) Evolutionary conservation of S326 and S330 in NADK2. D) The domain structure of NADK2, with serines 326 and 330 falling near the end of the kinase catalytic domain. E) Overview of RNAseq and metabolomics studies. Samples from *Nadk2* and *Pla2g6* mutant mice, as well as wild-type littermates for each, were collected at both early and late time points as described in the Methods. At each time point, samples were collected for RNAseq and metabolomics, as well as for histology. Samples for RNAseq derived from whole brain and muscle, and were sequenced using 100-bp paired-end sequencing. Samples for metabolomics included whole brain, muscle, spinal cord, and liver, as well as isolated mitochondrial preparations from brain and liver. These samples were analyzed by general GCTOF mass spectrometry, HILIC for biogenic amines, and lipidomics (positive and negative ionization modes).

The mouse NADK2 protein is 452 amino acids, encoded by the 14 exon *Nadk2* gene. The mutations identified are located at the 3’ end of exon 10 and the 5’ end of exon 11 (transcript *Nadk2*-205). Alternatively-spliced transcripts lacking either exon 5 or exon 8 are reported in the Ensembl genome browser (transcripts *Nadk2*-204 and −206), but exons 10 and 11 are included in all mRNAs producing full length protein. The two mutated serine residues at positions 326 and 330 are conserved through vertebrate evolution, and S326 is also conserved in *Caenorhabditis elegans* and *Drosophila melanogaster* (Figure 1C). The serine residues fall at the C-terminal end of the NAD kinase domain (Figure 1D). The mouse NADK2 protein is 452 amino acids, whereas the human protein is 442 amino acids, but the domain structure and the mutated serines are conserved.

To better understand the pathogenic mechanisms that lead to the *Nadk2* phenotype, we performed both metabolomic and transcriptomic analyses (Figure 1E), after confirming that the *Nadk2^-/-^* mice do indeed have neuromuscular and neurodegenerative phenotypes by histology. Furthermore, we compared the *Nadk2^-/-^* mice to a mouse mutation in *Pla2g6* (*Pla2g6^m1J^*, MGI:4412026, homozygotes hereafter referred to as *Pla2g6^-/-^* for simplicity), a model of infantile neuroaxonal dystrophy (15). The *Pla2g6*^-/-^ mice also have a neuromuscular phenotype that is outwardly similar to the *Nadk2^-/-^* mice, but through an independent gene and possibly unrelated pathogenic mechanism. We compared these genotypes to gain a better understanding of both *Nadk2* and *Pla2g6* mutation pathogenicity, to identify metabolomic and transcriptomic signatures that may be shared as general neuromuscular disease perturbations, and to identify genomic and metabolomic changes that are genotype-specific.

### Neuromuscular and neurodegenerative phenotypes in *Nadk2^-/-^* mice

The *Nadk2^-/-^* mice have overt neuromuscular phenotypes, including muscle wasting, tremor, abnormal gait, and hindlimb dysfunction. To better understand the basis for this, we performed histopathology on 3 males and 3 females at 5-weeks and 11-weeks-of-age, comparing to wildtype littermates. These ages represent an early time point, when overt phenotypes are just discernable, and a late-stage time point, when mice are clearly affected and may soon die. Consistent with the overt phenotype, cross sections of the gastrocnemius muscle showed striking progressive abnormalities (Figure 2 A-C). At five-weeks-of-age, occasional central nuclei were seen, indicating regenerated muscle fibers, and some atrophied fibers were present, but defects were subtle (Figure 2B). By eleven-weeks-of-age, muscle was severely atrophied, and most fibers were smaller than wild type (Figure 2C). This atrophy may be largely neurogenic, secondary to denervation, but myogenic defects may also be involved, as indicated by our-omics analyses below. Neuromuscular junctions in the hindlimb showed marked denervation, with many postsynaptic sites being completely devoid of overlying motor nerve terminals (Figure 2 D-F). The generally well-preserved integrity of the denervated postsynaptic sites at five-weeks-of-age is consistent with recent denervation (Figure 2E), before the postsynaptic apparatus has disintegrated and before the muscle fiber has atrophied. In the motor branch of the femoral nerve, which innervates the quadriceps muscle in the hind limb, myelinated axons were not reduced in number, but were reduced in size by eleven-weeks-of-age (Figure 2 G-L). The mutant axons were not smaller at five-weeks-of-age, and while the wild type axons significantly increased their size between 5 and 11 weeks, the mutant axons did not.

**Figure 2.**
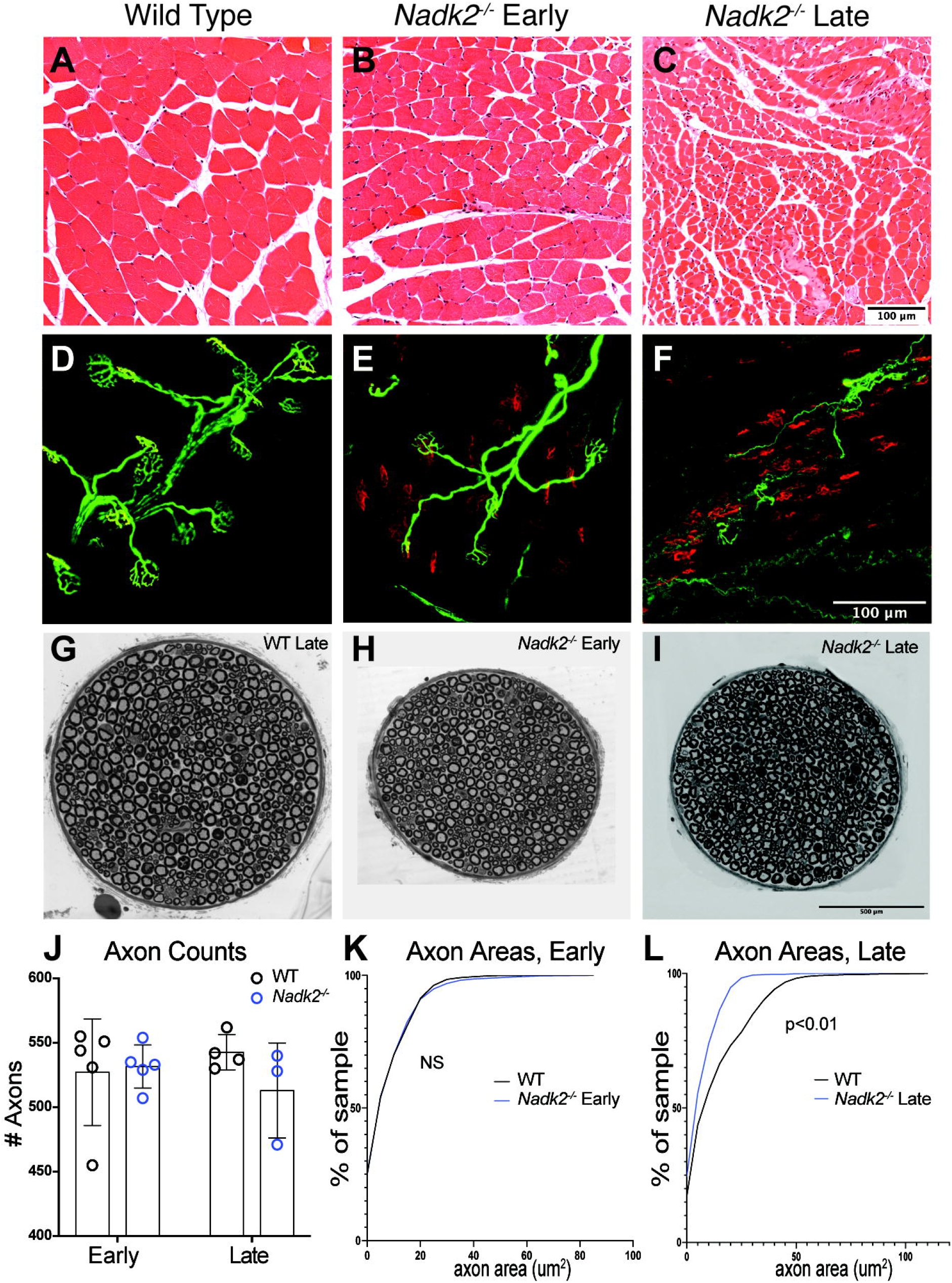
*Nadk2^-/-^* muscle and nerve histology. A-C) Cross sections of the gastrocnemius muscle stained with H&E in wild type (A), 5-week-old *Nadk2^-/-^* (B) and 11-week-old *Nadk2^-/-^* mice (C). Severe muscle atrophy was present at 11 weeks. D-F) Neuromuscular junctions in the plantaris muscle in wild type (D), 5-week-old *Nadk2^-/-^* (E) and 9-week-old *Nadk2^-/-^* mice (F). Neurofilament and SV2 label the nerve (green) and a-Bungarotoxin labels the postsynaptic acetylcholine receptors (red). Many denervated NMJs are observed at both 5- and 11-weeks-of-age. G-I) Cross sections of the motor branch of the femoral nerve in 11-week-old wild-type (G), 5-week-old *Nadk2^-/-^* (H) and 11-week-old *Nadk2^-/-^* mice (I). J) The number of myelinated axons in the femoral motor nerve was not reduced at either age in *Nadk2^-/-^* mice. K-L) The distribution of axon cross-sectional diameters was not different than control at 5-weeks-of-age (K), but axon size did not increase with age and axons in *Nadk2^-/-^* mice were smaller than wild-type at 11-weeks-of-age. Scale bar in C is 100 μm for A-C, in F is 100 μm for D-F, and in I is 500 μm for G-I. Significance tested by Student’s t-test (J), or Kologmorov-Smirnoff and Mann-Whitney U tests (K,L).

In addition to the motor axon loss, denervation, and muscle atrophy described above, we also observed neurodegeneration in other regions of the central nervous system in *Nadk2^-/-^* mice. Since optic neuropathy was described in at least one *NADK2* patient, we examined the retina in *Nadk2^-/-^* mice. There was no evidence of optic neuropathy, with no cupping at the optic nerve head (Figure 3 A-C), and retinal ganglion cell density was not reduced (14.2 ± 1.6 cells per 100 μm in control versus 15.2 ± 0.8 in 11-week *Nadk2^-/-^* mice). The outer nuclear layer of photoreceptors appeared less dense in mutant samples, but quantification revealed no changes in outer nuclear layer thickness or photoreceptor cell density (Figure 3D). In the cerebellum, there was progressive Purkinje cell loss, with a significant reduction in Purkinje cell density at 5-weeks-of-age, which was even more pronounced at 11-weeks-of-age (Figure 3E-H). An examination of sagittal sections of the brain did not reveal evidence of widespread neurodegeneration (pyknosis, chromatolysis, karyorrhexis) or astrocyte gliosis in regions such as the hippocampus or cortex (not shown). However, while our data above indicate that the *Nadk2* mutant mice have aspects of neurodegeneration, we acknowledge that this is not an exhaustive analysis and other cell populations maybe undergo neurodegeneration that we did not note with this high-level analysis.

**Figure 3.**
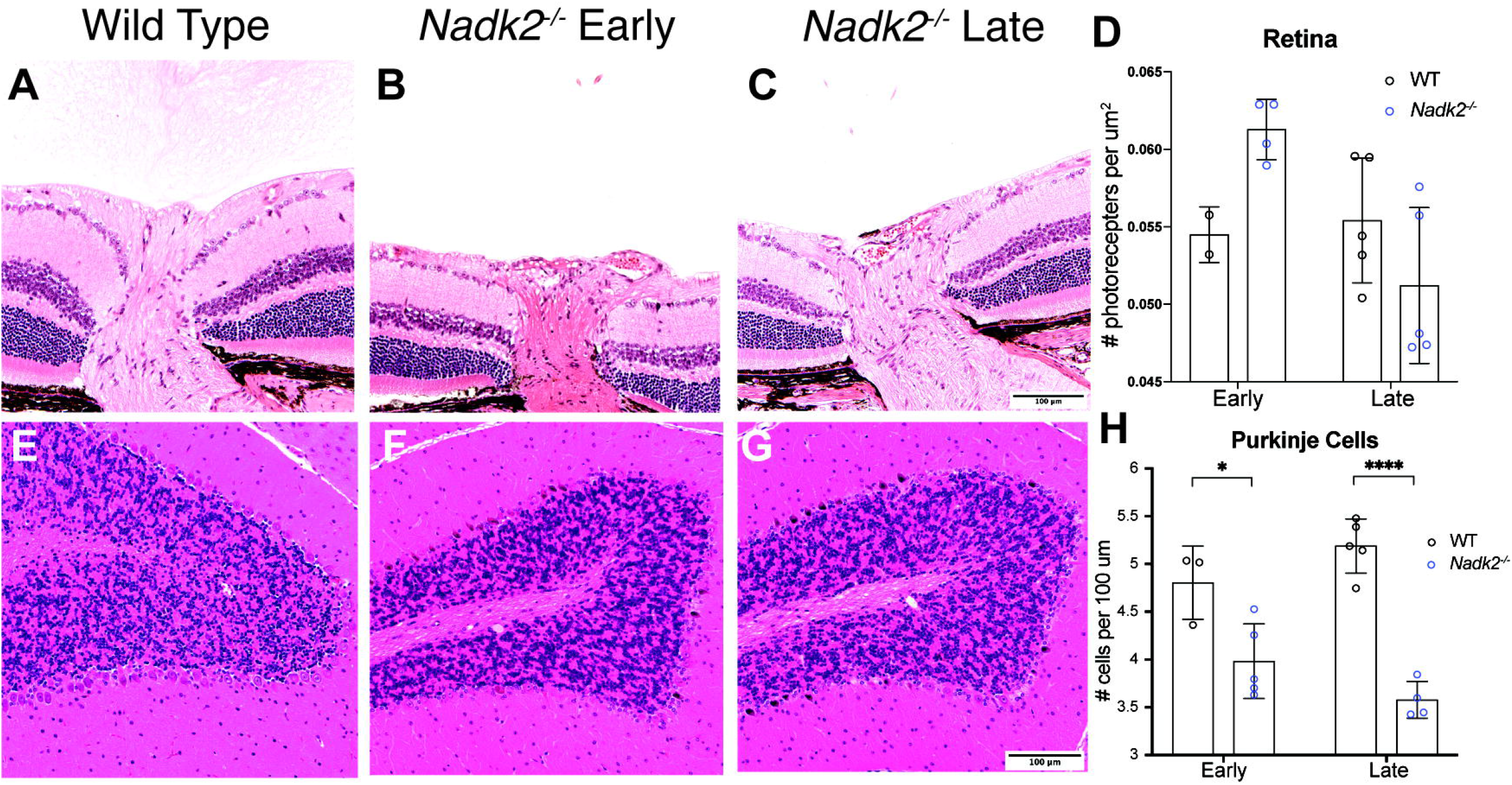
*Nadk2^-/-^* retina and cerebellum histology. A-C) Cross sections of the retina and the optic nerve head for wild type (A), 5-week-old (B), and 11-week-old (C) *Nadk2^-/-^* mice. No optic nerve cupping indicative of optic neuropathy was observed. D-F) Sagittal sections of the cerebellum in wild type (D), 5-week-old (E), and 11-week-old (F) *Nadk2^-/-^* mice. G) Quantification of photoreceptor number did not reveal a decrease in the *Nadk2^-/-^* mice at either early or late time points. H) Quantification of Purkinje cell number revealed a decrease in 5-week-old *Nadk2^-/-^* mice that was even more pronounced at 11-weeks-of-age. * p<0.05, **** p<0.0001. Pairwise comparisons tested using Student’s t-test. Scale bar in C is 100μm for A-C, and in F is 100 μm for D-F.

### RNAseq

In order to gain insight into the pathological mechanisms of *NADK2* deficiency, we performed RNA sequencing (RNAseq) of a sagittal half brain and skeletal muscle (triceps surae) from *Nadk2^-/-^* and *Pla2g6^-/-^* mice, as well as littermate controls for each (Figure 1E, Table 1). Two timepoints were used, 5-weeks and 11-weeks for *Nadk2* mutants, and 6-weeks and 13-weeks for *Pla2g6* mutants. These ages were selected to be either early or late in disease progression, and to match those used for the histological analyses described above. To identify the tissues and timepoints that exhibit transcriptomic and metabolomic disease signatures that differentiate between mutant samples and wild-type controls, principal component analysis (PCA) was used (Fig. 4 A,B). These plots indicate that clear separation of mutant samples from controls was only obtained from muscle at the late timepoint in *Nadk2^-/-^* mice, suggesting that muscle may be more strongly affected than brain in these animals. To examine the magnitude of the changes to the transcriptome and the similarity between data sets, we first generated lists of differentially expressed genes by comparing mutant to control within each time point and tissue type with cutoffs for significance and fold change (FDR ≤ 0.05 and |FC| ≥1.5). These lists were then compared as shown in the Venn diagrams (Figure 4C). These diagrams suggest that the disease signature seen in muscle from late *Nadk2^-/-^* mice is already at least partly present in the early timepoint, given the overlap in the gene lists. In contrast, the number of differentially expressed genes is smaller and there is little overlap between muscle samples from early and late *Pla2g6^-/-^* mice, suggesting that this mutation has a milder effect on the transcriptome of muscle than does *Nadk2^-/-^*. When differentially expressed genes are compared between skeletal muscle of *Nadk2^-/-^* and *Pla2g6^-/-^* mice at the later timepoint, it can be seen that, although there are many fewer differentially expressed genes in *Pla2g6^-/-^*, more than 50% of these differentially expressed genes are also seen in *Nadk2^-/-^*. This shared subset potentially includes genes that are generally misregulated in neuromuscular disease, while the majority of differentially expressed genes in *Nadk2^-/-^* are private and may represent pathways specific to that disease process. Using our stringent thresholds, many fewer differentially expressed genes were found in brain, and while early and late time points for each mutation overlapped, no differentially expressed genes were shared between *Nadk2^-/-^* and *Pla2g6^-/-^* samples (Figure 4D). We determined the number of differentially expressed genes from each mutant, tissue, and timepoint that have annotations in each of the databases we use for subsequent enrichment analysis (Supp Table 1).

**Figure 4.**
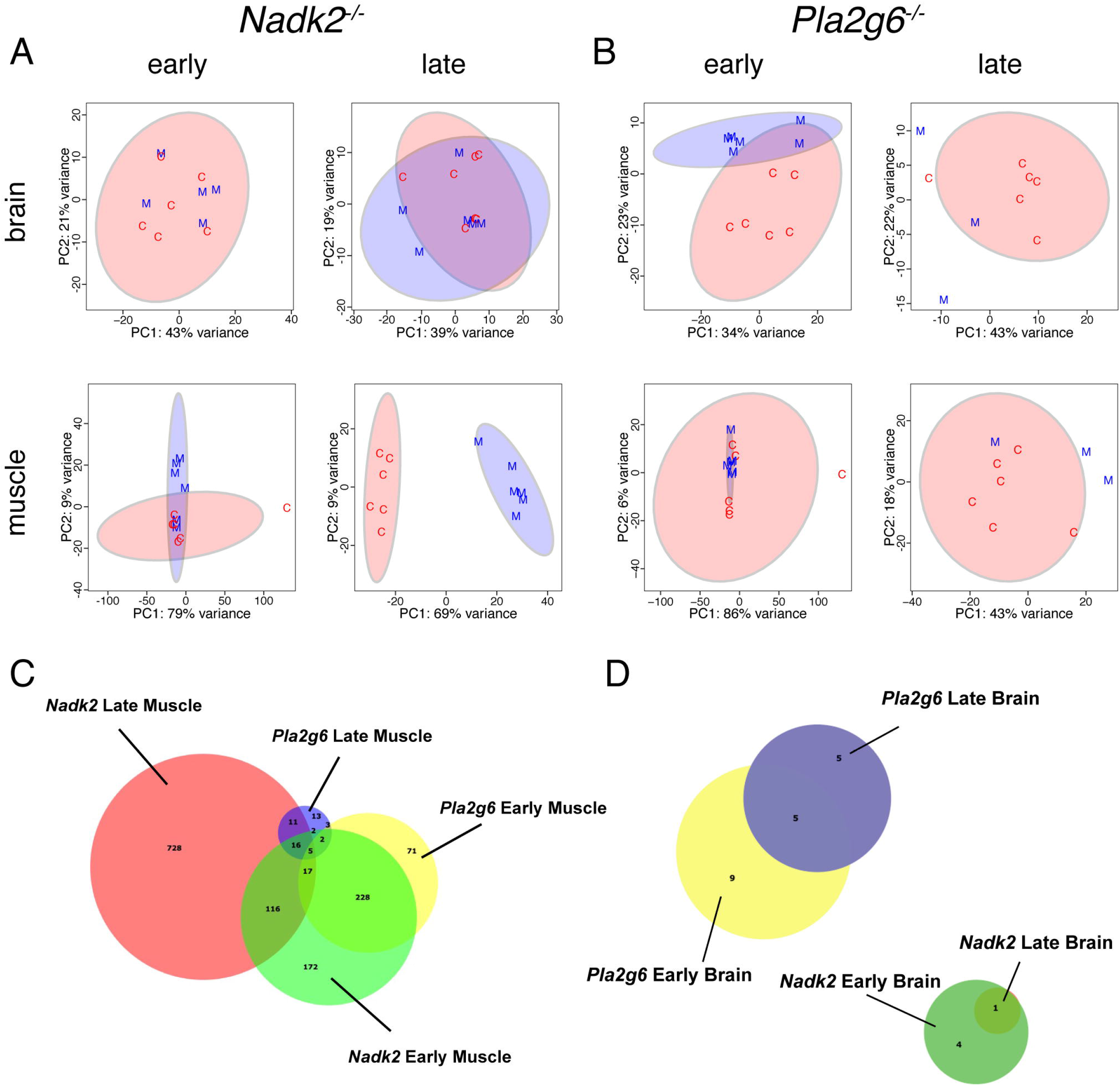
PCA plots and differential expression analysis of the transcriptome of *Nadk2^-/-^* and *Pla2g6^-/-^* mice. A) PCA plots of differentially expressed genes from brain and muscle of *Nadk2^-/-^* mice (M, blue) compared to wild-type littermate controls (C, red) at early and late timepoints. Clear separation of the genotypes is seen only at the late time point in muscle. B) PCA plots of differentially expressed genes from brain and muscle of *Pla2g6^-/-^* mice compared to wild-type littermate controls at early and late timepoints. Plots indicate little separation of the transcriptomes from the two groups. C, D) Venn diagrams comparing the number of differentially expressed genes and their overlap between the indicated genotypes and time points for muscle (C), and brain (D).

**Table 1:** RNAseq samples. Genotypes, ages, sex, and tissues used in RNAseq studies.

| Genotype | Age | N | Tissues |
| --- | --- | --- | --- |
| *Nadk2*<sup>-/-</sup> | 5 weeks | 3F, 3M | Brain<br>Muscle |
|  | 11 weeks | 3F, 3M |  |
| *Pla2g6*<sup>-/-</sup> | 6 weeks | 2F, 3M |  |
|  | 13 weeks | 2F, 2M |  |
| Wild type | 5-6 weeks | 3F, 3M |  |
|  | 11-13 weeks | 3F, 3M |  |

### Metabolomics

To complement the RNAseq data, samples were obtained from sagittal hemi-brains and muscle at early and late timepoints, as well as from spinal cord and liver, from both *Nadk2^-/-^* and *Pla2g6^-/-^* mice and littermate controls for metabolomics studies. Mitochondrial preparations from brain and liver from the same time points were also included (Figure 1E, Table 2). All samples were analyzed by GC-TOF mass spectrometry to screen for primary metabolites, and brain and muscle whole-tissue samples were additionally analyzed for biogenic amines by HILIC and for lipidomics by CSH-QTOF-MS/MS. Several approaches were taken to analyze the extent to which mutant samples differed from matched wild-type controls. First, Random Forest analysis was performed on each group, and the Out of Bag (OOB) errors were determined (Figure 5A). As lower OOB scores indicate better separation of mutant from control, *Nadk2^-/-^* overall showed a greater degree of difference from wild-type than did *Pla2g6^-/-^*. Specifically, samples from the late timepoint showed a greater difference from wild-type than did samples from the early timepoint, and samples from muscle showed the greatest degree of difference from wild-type. Samples from late *Nadk2^-/-^* muscle yielded OOB error scores of 0 on all 3 metabolomics platforms utilized, suggesting a large magnitude of change within the metabolome of muscle by the late stages of disease. The degree of separation between select *Nadk2^-/-^* and wild-type samples was also visualized by plotting scores from Principal Component Analysis (PCA; Figure 5 B,C). In agreement with the Random Forest OOB scores, clear separation of mutant and control samples was only seen in *Nadk2^-/-^* muscle at the late timepoint, and this separation was seen using data from any of the 3 metabolomics platforms. PCA was not performed on *Pla2g6*^-/-^ samples, given the comparatively small number of differentially detected metabolites. Thus, our metabolomics results agree with the major conclusion of the RNAseq described above, that the strongest differences are seen in *Nadk2^-/-^* muscle at the late timepoint.

**Figure 5.**
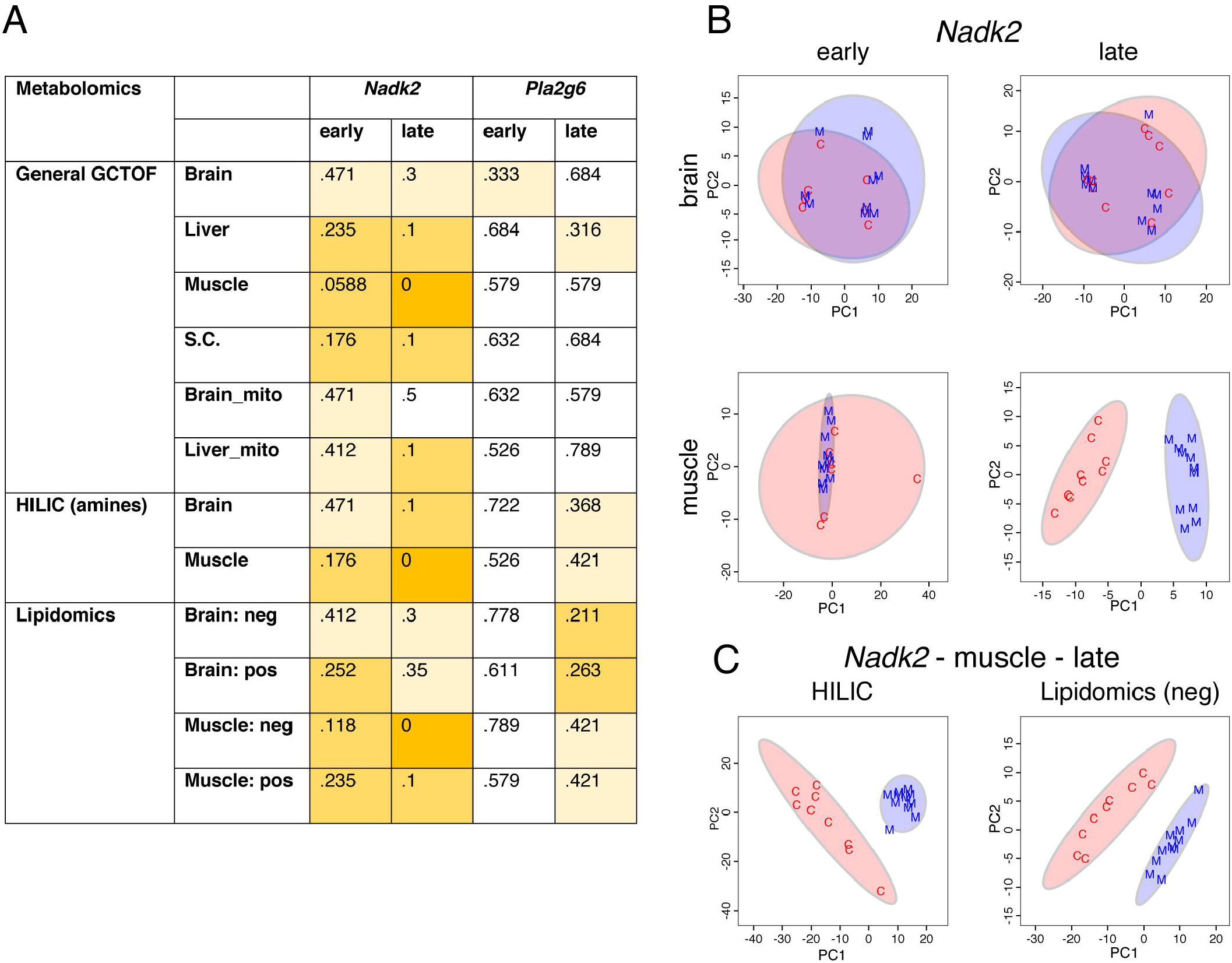
Random Forest Out of Bag (OOB) error rates and PCA scores plots for metabolomics analysis of *Nadk2* and *Pla2g6* mutant mice versus littermate controls. A) OOB error rates from Random Forest analysis of all metabolomics data from mutant mice compared to littermate controls. Scores are color-coded to indicate the magnitude of the error score, with scores of zero shaded dark orange, those between zero and 0.3 shaded medium orange, and those between 0.3 and 0.5 shaded pale orange. The lowest error rates are seen in *Nadk2^-/-^* mice at the late time point, especially in muscle samples. B) PCA scores plots of general GCTOF metabolomics data of *Nadk2^-/-^* mice (M, blue) versus wild-type littermate controls (C, red) from brain and muscle samples at early and late time points. Separation of the genotype groups is only observed at the late time point in muscle. C) PCA scores plots of HILIC and lipidomics analyses from muscle collected from *Nadk2^-/-^* mice at the late time point also display separation from matched wild-type samples.

**Table 2:** Metabolomics samples. Genotypes, ages, sex, and tissues used in metabolomics studies.

| Genotype | Age | N | Tissues |
| --- | --- | --- | --- |
| *Nadk2*<sup>-/-</sup> | 5 weeks | 4F, 6M | Brain<br>Muscle<br>Spinal Cord<br>Liver<br>Brain-mito<br>Liver-mito |
|  | 11 weeks | 5F, 5M |  |
| *Nadk2*<sup>+/+</sup> | 5 weeks | 2F, 5M |  |
|  | 11 weeks | 3F, 6M |  |
| *Pla2g6*<sup>-/-</sup> | 6 weeks | 5F, 4M |  |
|  | 13 weeks | 4F, 6M |  |
| *Pla2g6*<sup>+/+</sup> | 6 weeks | 2F, 7M |  |
|  | 13 weeks | 4F, 5M |  |

### Integrated Transcriptomic and Metabolomic Analysis of *Nadk2* deficient mice

We next integrated the results of brain and muscle RNASeq and primary metabolite levels on the GCTOF platform from *Nadk2^-/-^* mice at 5 and 11 weeks of age. At the late 11-week timepoint lysine levels were significantly higher in *Nadk2^-/-^* muscle tissue than wild-type controls (Figure 6A). Lysine was also significantly elevated at the early 5-week timepoint in *Nadk2^-/-^* muscle, whereas lysine levels were not significantly elevated in *Nadk2^-/-^* brain, despite a strong trend (Supp Figure 1A). Proline is also significantly elevated in late muscle tissue of *Nadk2^-/-^* mice, and glucose-derivative sorbitol is significantly reduced (Figure 6A), reflecting the dependence of enzymes in the polyol pathway on NADP as a cofactor. There are 918 differentially expressed genes (DEGs) in *Nadk2^-/-^* muscle relative to wild-type controls at the late timepoint, which is more than any other tissue or timepoint for *Nadk2^-/-^* mice (Figure 4C and Figure 6B). Of the 557 DEGs present in early *Nadk2^-/-^* muscle, 154 are present in both early and late muscle. There are only five DEGs in brain at the early timepoint, and only one at the later timepoint (Supp Figure 1B). At the late timepoint, DEGs in *Nadk2^-/-^* muscle are associated with Mammalian Phenotype Ontology (MPO) terms: skeletal muscle, the nervous system, body mass, and metabolism. Reactome pathway associations include muscle contraction and postsynaptic acetylcholine receptors (Figure 6C). DEGs unique to the early timepoint in muscle have highly significant associations with MPO terms related to central nervous system structure and function. Those DEGs that overlap between early and late muscle are associated with body mass, and those DEGs unique to the late timepoint are associated with behavior and physiological MPO terms (Supp Figure 1C).

**Figure 6.**
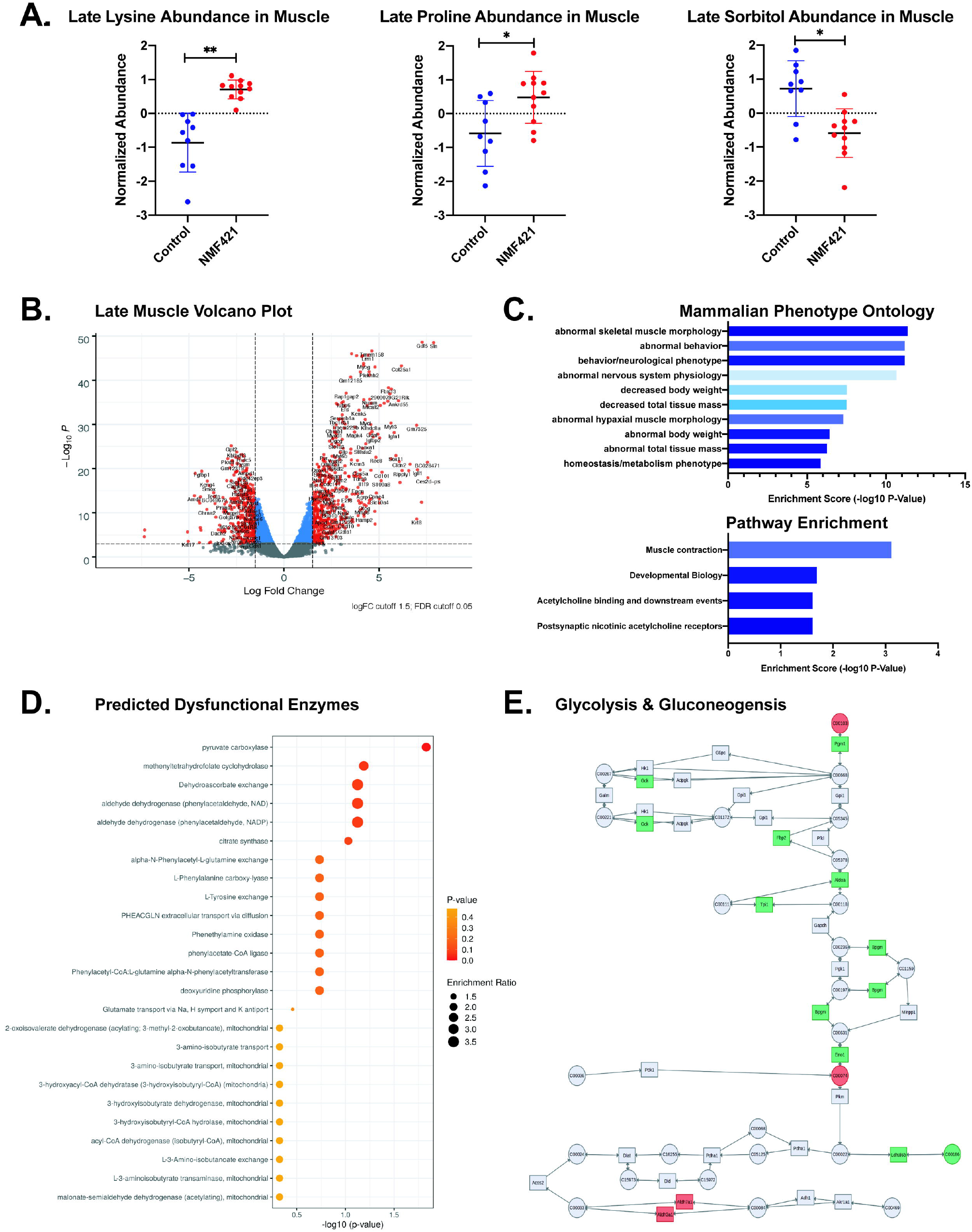
Transcriptomic and Metabolomic Analysis of *Nadk2^-/-^* Muscle. A) The normalized abundance of lysine, proline, and sorbitol by sample in muscle tissue of *Nadk2^-/-^* mice at 11 weeks of age. Significance is determined by MetaboAnalyst5.0 t-test. B) Volcano plot of differentially expressed genes in *Nadk2^-/-^* muscle at the late timepoint. Genes depicted in red passed filtering and were used for ontology searches and downstream analysis. C) Mammalian Phenotype Ontology and pathway ontology terms associated with differentially expressed genes ranked by p-value. D) MetaboAnalyst5.0 predicted dysfunctional enzymes based on significant differences in metabolite abundance in *Nadk2^-/-^* muscle at 11 weeks. Color coded to represent p-value and size scaled based on enrichment ratio. E) Joint pathway analysis in MetaboAnalyst5.0 identifies glycolysis & gluconeogenesis as the top dysfunctional pathway to pass FDR ≤ 0.05 cutoff ranked by impact score. Color coded to indicate direction of differential expression, red indicates high abundance or expression, and green represents decreased abundance of expression.

Metabolite enrichment analysis in MetaboAnalyst5.0 using metabolite lists from *Nadk2^-/-^* muscle at the late timepoint enabled prediction of enzymes whose deficiency would produce metabolomic changes consistent with uploaded lists (Figure 6D). The top predicted enzyme by significance is pyruvate carboxylase. In *Nadk2^-/-^* muscle at the early timepoint, “hyperlysinemia I or saccharopinuria” is the top metabolite-disease association and has the largest enrichment ratio (Supp Figure 1D). 2,4-Dienoyl-CoA Reductase Deficiency is also present in this list, along with a variety of other metabolic deficiencies and nervous system disorders. When DEGs were uploaded alongside metabolite lists in MetaboAnalyst5.0 Joint-Pathway module the top predicted pathway by impact with FDR ≤ 0.05 was glycolysis and gluconeogenesis (Figure 6E and Supp Figure 2A). We also performed a targeted search for *NADK2* deficiency associated pathways “lysine degradation” and “beta oxidation” in the KEGG pathway database and used the mapper tool to project DEGs onto these pathways (Supp Figure 2B, C). In both cases, numerous DEGs from our lists were involved in target pathways (Beta oxidation: M00087: 40/55, Lysine degradation: M00032: 4/10). Gene set enrichment analysis (GSEA) of Trimmed mean of M-value (TMM)-normalized counts from *Nadk2^-/-^* muscle at the late timepoint identified enrichment of a diverse set of Reactome pathways, most of which are related to metabolism. Two relevant pathways with high magnitude normalized enrichment scores when *Nadk2^-/-^* mutants are compared to controls in GSEA were Mitochondrial Fatty Acid Beta Oxidation and Pyruvate Metabolism, and Citric Acid TCA Cycle (Supp Figure 1E), which indicates metabolic disruption is the most prominent ‘omics feature across multiple analysis methods.

### Transcriptomic Analysis of *Pla2g6^-/-^* mice

Since no metabolites from *Pla2g6^-/-^* mice had significant differences in abundance after correction for multiple testing (FDR ≤ 0.05), we focused our analysis on differentially expressed genes (DEGs). There are 14 DEGs in *Pla2g6*^-/-^ brain at the early timepoint, and 10 at the late timepoint. There are 349 DEGs in *Pla2g6*^-/-^ muscle at the early timepoint, and 52 DEGs at the later timepoint. Only three DEGs overlap between both tissues and both timepoints, and one of these is *Pla2g6* (Supp Figure 3A, B). The other two DEGs encode ribosomal proteins (Supp Figure 2B). Given the paucity of DEGs in *Pla2g6*^-/-^ brain we focused our analysis on muscle. In general, DEGs in early tissue exhibited higher magnitude fold changes at approximately equal significance levels, and more genes were down-regulated than up-regulated in early samples (Supp Figure 3C, D). DEGs from early muscle tissue were associated with Reactome pathways related to intracellular signaling cascades and MPO terms related to the nervous system and especially synaptic transmission (Supp Figure 3E). DEGs at the late timepoint are associated with only two Reactome pathways: striated muscle contraction and muscle contraction. MPO terms relate to the heart and cardiac function (Supp Figure 3F). Though no disease-associated terms reached significance after correction for multiple testing, “neurodegeneration with brain iron accumulation” had a nominal p-value < 0.05 at both early and late timepoints in brain (Supp Figure 3G). It should be noted that these associations arose with a relatively small number of annotated genes (Supp Table 1).

### Comparative Analysis of *Nadk2* and *Pla2g6* mutant mice

To better understand the transcriptomic changes specific to *Nadk2* or *Pla2g6* mutants, and those changes that might overlap between neuromuscular disorders, we examined the overlap between DEGs of each tissue, genotype and timepoint. Only DEGs from muscle overlapped between mutants, and this overlap was greatest at the early timepoint (Figure 7A). Overlap between enriched Reactome pathways in GSEA with FWER < 0.05 was also greatest in muscle at the early timepoint (Figure 7B). There are 32 genes that overlap between *Nadk2* and *Pla2g6* mice at the late timepoint. Only gene ontologies (GO:BP) related to muscle development are significantly associated with this late gene set. Reactome pathways associated with the 251 DEGs shared between *Nadk2^-/-^* and *Pla2g6*^-/-^ muscle at the early timepoint are associated with the nervous system and include numerous calcium-dependent processes and neurotransmitter releases processes by over-representation analysis (Figure 7C). Gene set enrichment analysis (GSEA) identified a wider array of Reactome pathways. Top pathways with a positive normalized enrichment score (NES), i.e. those pathways associated with positive log2FC genes in mutants versus controls, are associated with translation, while those with a negative NES, associated with negative log2FC genes, are related to synaptic signaling (Figure 7D).

**Figure 7.**
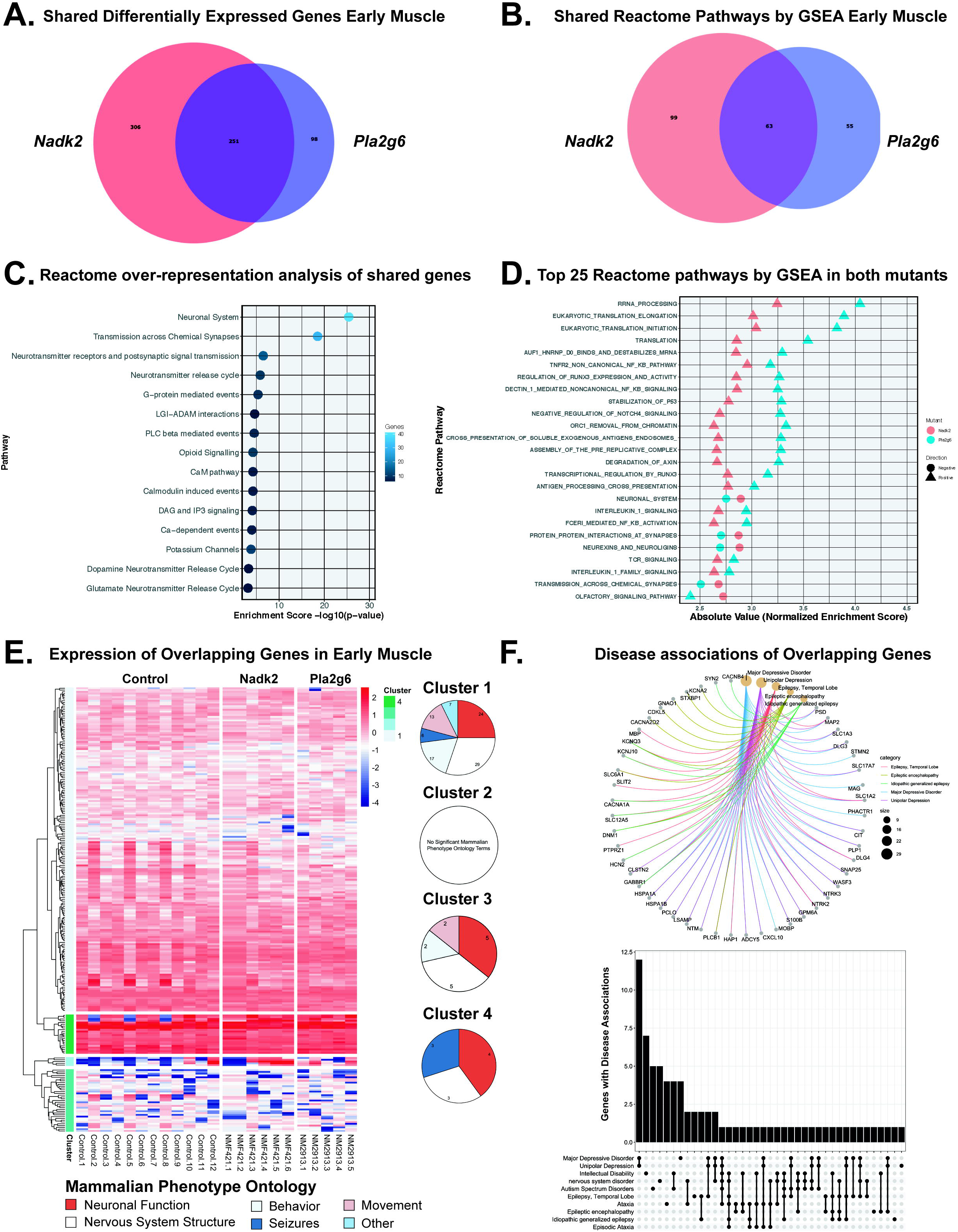
Comparative transcriptomic analysis of *Nadk2* and *Pla2g6* mutants. A) Venn diagrams depicting overlapping differentially expressed genes in *Nadk2^-/-^* and *Pla2g6^-/-^* muscle at the late timepoints. B) Venn diagram depicting overlapping Reactome pathways implicated by Gene Set Enrichment Analysis with FWER < 0.05. C) Pathway ontology terms associated with differentially expressed genes shared between *Nadk2^-/-^* and *Pla2g6^-/-^* muscle at the early timepoint. Color indicates the number of genes associated with each ontology term. D) Top 25 Reactome pathways by Normalized Enrichment Score with FWER < 0.05 implicated by Gene Set Enrichment Analysis shared between *Nadk2^-/-^* and *Pla2g6^-/-^* muscle at the early timepoint. Color indicates mutant and shape differentiates positive and negative normalized enrichment scores. E) Heatmap and dendrogram depicting expression level of differentially expressed genes shared between *Nadk2^-/-^* and *Pla2g6^-/-^* early muscle tissue. Genes are clustered into four groups based on the similarity of gene expression patterns across samples. Ontology terms for each cluster are grouped into major categories, and the representation of major categories for each cluster are presented as circular plots. F) Circular association and upset plots of gene-disease associations from human orthologs of differentially expressed genes that overlap between *Pla2g6^-/-^* and *Nadk2^-/-^* early muscle.

To determine if the direction of differential expression was consistent between mutants, we created a heatmap using TMM-normalized counts. We applied a supervised clustering approach based on dendrogram height to segment these genes into four clusters based on expression: genes with moderate expression across all samples (cluster 1), genes with very high or very low expression that vary widely between samples (cluster 2), genes with moderate expression that vary widely across samples (cluster 3), and genes with high expression across all samples (cluster 4). We examined MPO terms separately for each cluster and determined that cluster 4 genes were disproportionately associated with neuronal functions and seizures, while cluster 1 genes have diverse associations with more movement- and behavior-related terms than other clusters (Figure 7E). We examined the gene-disease associations of human orthologs of DEGs shared between *Nadk2* and *Pla2g6* mutant muscle at the early timepoint. Top disease associations were limited to disorders of the central nervous system, but also include ataxia (Figure 7F).

## Discussion

Here we describe two mutations in mouse *Nadk2* that were identified because of their overt neuromuscular symptoms. Both mutations alter serine residues near the C-terminal end of the catalytic domain of NADK2 and are likely partial loss-of-function alleles. Histological examination of *Nadk2^-/-^* mice indicates there are indeed neuromuscular and neurodegeneration phenotypes, with muscle atrophy and denervation, as well as loss of cerebellar Purkinje cells. Analysis of both differential metabolites and differentially expressed genes in several tissues demonstrates that changes in muscle best distinguish mutant mice from controls. The pathways perturbed in *Nadk2^-/-^* mice were largely separate from those affected in *Pla2g6^-/-^* mice, despite the similar overt presentation of the mutant animals. An integrated analysis of altered genes and metabolites demonstrates that the *Nadk2^-/-^* mice recapitulate major hallmarks of the patients, including hyperlysinemia and deficiencies in beta-oxidation of fatty acids, mediated partially by DECR1. Other pathways such as the polyol pathway, which has been recently linked to peripheral neuropathy (17), were also altered.

Although a limited number of *NADK2* patients have been described, they consistently present with neurological and metabolic conditions, including hyperlysinemia and DECR-deficiency. The identification of similar symptoms in the *Nadk2^-/-^* mice suggest that the S330P mutation creates a valid model of the human disease. This mutation is likely a hypomorph, as only a single amino acid is changed and the phenotype is milder than null mice produced through the KOMP program, which die embryonically. However, the mice have a more severe phenotype than described for the *Nadk2*^tm1a(EUCOMM)Wtsi^ allele, which may be an even weaker hypomorph with reduced levels of wild type *Nadk2* produced. The relevance of our findings in this mouse model to the human disease are discussed below.

Our genetic analysis also indicates that serines 326 and 330 are both important for NADK2 function, as mutations at either of these residues produces a strong partial-loss-of-function phenotype. We have admittedly not explored the impact of these mutations on NADK2 enzymatic activity in this study. These mutations could have multiple (and not mutually exclusive) possible impacts on NADK2, including its enzymatic activity (kinetics, substrate specificity, etc.) or its regulation through modifications such as through phosphorylation or other post-translational modifications. The serine at 326 is predicted to be a possible protein kinase C (PKC) target using online tools, but this has not been experimentally verified. These studies would best be done by investigators with strong expertise in kinase biochemistry.

The hyperlysinemia and C10:2 carnitine elevation may be understood through the requirement for NADP as a substrate for AASS and DECR1 in lysine degradation and beta-oxidation of fatty acids, respectively. However, the severe disease phenotype and neurological symptoms are likely not wholly attributable to these pathways. To characterize the broader cellular dysfunction underlying neuromuscular deficits in our *Nadk2^-/-^* mice we performed transcriptomic and metabolomic analyses at the level of individual genes and metabolites, ontologies associated with gene and metabolite lists, and pathway analysis that integrated both differentially expressed genes and metabolites. Our *Nadk2^-/-^* mice exhibited significant increases in lysine abundance in muscle tissue at both early and late timepoints. At the early timepoint, the metabolite list from muscle showed very high enrichment ratios against hyperlysinemia and 2,4-dienoyl-CoA reductase deficiency metabolites sets. At the later timepoint, the metabolite sets by enrichment ratio pertained to dysfunction of metabolic enzymes such as pyruvate carboxylase. However, mapping differentially expressed genes names onto *NADK2* deficiency KEGG pathways “lysine degradation” and “beta-oxidation of fatty acids” revealed numerous overlaps at the later timepoint (7). This suggests that early onset deficiencies in key pathways are later masked by a progressive generalized metabolic dysfunction in ‘omics data, but are still present. Mammalian Phenotype Ontology terms from differentially expressed genes in muscle tissue pertain to the nervous system at an early timepoint but transition almost entirely to terms associated with abnormalities of skeletal muscle at the later timepoint. This finding parallels the denervation of neuromuscular junctions and muscle atrophy identified by histology. The muscle atrophy in these mice is progressive and likely neurogenic. We also conclude that our *Nadk2* mutant mice broadly recapitulate biochemical signatures of *NADK2* deficient patients.

We also analyzed a *Pla2g6* mouse model of INAD using similar methods. A major difference between *Nadk2* and *Pla2g6* mutant mice is the absence of metabolites that passed FDR ≤ 0.05 filtering in any tissue or timepoint of *Pla2g6^-/-^*. Analysis of differentially expressed genes in early timepoint muscle identified annotations involving the nervous system. These terms are replaced by annotations associated with muscle contraction, and heart muscle contraction, at the later timepoint. Very few ontology terms reached significance in brain tissue from *Pla2g6* mutants, and for this reason we considered terms that fell short of Holm-Bonferroni multiple testing correction, using a nominal p-value as an exploratory method. Interestingly, this approach revealed a small number of highly specific terms such as “neurodegeneration with brain iron accumulation”, likely driven by the reduced expression of *Pla2g6* itself. Muscular pathology has been documented in *PLA2G6* neuroaxonal dystrophy patients, but generally brain iron accumulation is considered a defining feature of this disorder (18,19). These results assured us that our analysis was sensitive enough to detect disease-relevant molecular signatures in both *Nadk2* and *Pla2g6* mutant mice. However, we were interested in determining if our identification of disease-relevant biochemical signatures had high specificity for each mutant. Given the absence of metabolites that passed FDR filtering for *Pla2g6^-/-^*, we compared lists of differentially expressed genes. No differentially expressed genes overlapped between brain of the two mutants, and so our comparison was limited to muscle. At the later timepoint, a small number of Mammalian Phenotype Ontology terms related to skeletal muscle were significant. We then focused on the early timepoint where 251 differentially expressed genes are shared. Over-representation analysis suggested these shared genes were related to the nervous system and chemical synaptic transmission, perhaps driven by postsynaptic response to denervation. Gene Set Enrichment Analysis generally agreed with our over-representation analysis, but also identified shared positive enrichment of translation processes between mutants. Our cluster analysis indicated that neurological terms are consistently implicated by differentially expressed genes regardless of expression level with a few minor deviations noted earlier. Gene-disease associations implicate a general set of nervous system disorders. Our failure to identify annotation enrichment for *Nadk2*-related processes such as metabolism and lysine-degradation or *Pla2g6*-related processes such as brain iron accumulation and calcium-dependent events in the shared set suggests our analysis reliably discriminated pathways that are specific to each condition. However, our transcriptomic and metabolomic analyses probably do not fully reflect the underlying pathophysiology of each mutant with complete fidelity, given the tendency for annotation bias to skew results in favor of well-studied processes (20). Nonetheless, the gene expression and metabolomics datasets reported here will aid in the study of human *NADK2* mutations and infantile neuroaxonal dystrophies. The S330P *Nadk2* mutant mice may be better suited for *in vivo* studies than previous models of this rare recessively inherited condition.

## Methods

### Mapping and identification of *Nadk2* mutations

We identified two strains of mice with an overt neuromuscular phenotype (hind limb wasting, kyphosis, poor gait and motor ability) from two independent screens using N-ethyl-N-nitrosourea (ENU) to mutagenize mice. The screens have been previously described (21,22). Both mutations were assumed to be single-gene recessive events based on pedigrees.

The first allele, referred to as *m1Jcs*, was ENU induced in a C57BL/6J background. The mutation was mapped by transferring ovaries from an affected (presumed homozygous) mouse into NOD.CB17-*Prkdc*^scid^/J recipient female. This mouse was subsequently bred to CAST/Ei, producing F1 mice that were presumed obligate heterozygotes. F1 mice were intercrossed to produce F2 mapping animals. A genome scan and subsequent fine mapping showed complete linkage to proximal Chr. 15 between *D15Mit224* and *D15Mit11*. This region includes ten protein coding genes (*Capsl, Il7r, Lmbrd2, Nadk2, Ranbp3l, Skp2, Slc1a3, Spef2, Ugt3a1, and Ugt3a2*). The coding portion of the candidate genes in the interval were evaluated by Sanger sequencing of PCR amplified whole brain cDNA produced from mutant and heterozygote mapping animals.

The second allele, designated *nmf421*, was induced in a C57BL/6J background and was mapped by breeding an early-stage affected mouse (presumed homozygote) to BALB/cByJ to produce F1 mice (presumed obligate heterozygotes). The F1 offspring were then intercrossed to produce F2 mice used in mapping. A genome scan using simple sequence length polymorphisms (SSLPs, *Mit* markers) was performed and significant association with markers *D15Jmp22* (11.0 mb) and *D15Mit252* (22.7 mb) on proximal Chr. 15 was identified. Of 13 affected mice screened, 12 were homozygous C57BL/6J at both markers, one was heterozygous (Chi square test 2.3×10^-8^), and more distal markers on Chr. 15 were less-well associated, suggesting the mutation was proximal to *D15Jmp22*. Given the identification of the *m1Jcs* mutation in *Nadk2*, exons of *Nadk2* were sequenced in *nmf421*, identifying the *Nadk2^nmf421^* S330P allele.

#### Tissue collection

For all studies, *Nadk2* and *Pla2g6* homozygous mutant mice were produced in heterozygous matings, and littermates were used as controls (in rare cases, strain and age matched wild type mice were used to fill out a cohort). For metabolomics, mice were euthanized by cervical dislocation and organs were collected and snap frozen in liquid nitrogen as quickly as possible. One sagittal half brain and one lobe of the liver were used to prepare crude mitochondrial fractions by centrifugation, as described (23). Tissue was then powdered while frozen and separated into up to three sub-samples for extraction and mass spectrometry at the West Coast Metabolomics Center. Sample numbers, genotypes, ages and sexes used for metabolomics are given in Table 2. For RNAseq, a sagittal hemi-brain or triceps surae muscle were snap frozen in liquid nitrogen for subsequent RNA isolation. The other half brain and muscles from the other leg were processed for histology as described below. Tissue was collected in batches (litter by litter), and banked at −80°C. All samples were processed and analyzed (metabolomics or RNAseq) in single batches. Sample numbers, genotypes, ages and sexes for RNAseq are given in Table 1.

#### RNAseq

RNAseq was performed on the following samples: brain and muscle from *Nadk2* and *Pla2g6* mutants, along with wild-type littermates, at early (5wk for *Nadk2^-/-^* and 6wk for *Pla2g6^-/-^*) and late (11wk for *Nadk2^-/-^* and 13wk for *Pla2g6^-/-^*) timepoints. Tissue was collected, snap frozen in liquid nitrogen, and stored at −80 until it was processed for RNA extraction and library preparation. **RNA isolation:** RNA was isolated from tissue using the MagMAX mirVana Total RNA Isolation Kit (ThermoFisher) and the KingFisher Flex purification system (ThermoFisher). Tissues were lysed and homogenized in TRIzol Reagent (ThermoFisher). After the addition of chloroform, the RNA-containing aqueous layer was removed for RNA isolation according to the manufacturer’s protocol, beginning with the RNA bead binding step. RNA concentration and quality were assessed using the Nanodrop 2000 spectrophotometer (Thermo Scientific) and the RNA Total RNA Nano assay (Agilent Technologies). **Library preparation:** Libraries were prepared using the KAPA mRNA HyperPrep Kit (KAPA Biosystems), according to the manufacturer’s instructions. Briefly, the protocol entails isolation of polyA containing mRNA using oligo-dT magnetic beads, RNA fragmentation, first and second strand cDNA synthesis, ligation of Illumina-specific adapters containing a unique barcode sequence for each library, and PCR amplification. Libraries were checked for quality and concentration using the D5000 ScreenTape assay (Agilent Technologies) and quantitative PCR (KAPA Biosystems), according to the manufacturers’ instructions. **Sequencing:** Libraries were pooled and sequenced 70 bp single-end on the HiSeq 4000 (Illumina) using HiSeq 3000/4000 SBS Kit reagents (Illumina), targeting 30 million reads per sample.

Standard RNA-seq pipeline comprising of tools to perform read quality assessment, alignment, and variant calling was developed at The Jackson Laboratory. The pipeline takes sequence reads for each sample as raw fastq files and outputs read counts. Sequence base qualities ≥30 over 70% of read length criteria was used in downstream analyses. Sequence reads that passed the quality were aligned to a mouse reference (GRCm38.74) using Bowtie v2.2.0 (24,25). To ensure reliable quantification of gene expression level for all genes, ≥ 1 million mouse reads threshold was used for expression analysis. Gene expression estimates were determined using RSEM v1.2.19 (rsem-calculate-expression) with default parameters (26). Expression estimates were further normalized by using upper quantile normalization of non-zero expected counts and scaling to 1000. R package edgeR was used to perform the normalization and the test for differential expression (27). Normalization factors were calculated using TMM method (28). Constrained Regression Recalibration (ConReg-R) was used to recalibrate the empirical p-values by modeling their distribution to improve the FDR estimates (29). Gene lists were filtered using cutoffs for significance and fold change (FDR≤0.05 and |FC| ≥1.5). Raw data were deposited into GEO, accession number 188592, and raw differential expression data are included as Supplemental Data.

#### Metabolomics

Metabolomics analysis was performed at the West Coast Metabolomics Center. Three platforms were used. All samples (hemi-brain, muscle, liver and spinal cord, as well as mitochondria isolated from brain and liver) were examined for primary metabolism by GC-TOF-MS. Brain and muscle were also analyzed for lipidomics using CSH-QTOF-MS/MS (positive and negative electrospray ionization) and for biogenic amines by HILIC-QTOF-MS/MS.

Detailed methods provided by the WCMC are provided in the supplemental materials. The ages, genotypes, sex, and numbers of the mice analyzed are provided in Table 2. Concentration data were uploaded as comma separated values (.csv) files to MetaboAnalyst. Data integrity check, missing value imputations, and data filtering were performed before normalization. Statistical analysis was performed with the null hypothesis of no difference between the mutant and control samples at each time point. Multiple exploratory, univariate and multivariate, and machine learning methods were used in the data analysis (30,31). Samples with 50% or more values missing in either mutant or control or both samples and were discarded from further analysis. Missing values were then imputed using half the detection limit and filtered using inter quartile range (IQR) method (32,33). The data was then log2 transformed and auto scaled (34,35). Student’s *t*-test, Fold change, principal component analysis (PCA) were performed as exploratory data analysis methods. Machine learning models were developed using support vector machine (SVM) algorithm (36), with linear kernel method and Random Forest analysis (37) to classify the mutant and control samples and rank the importance of metabolites in discriminating the two types of samples (see Metabolomics Analysis Supplement for full details of data processing and analysis).

#### Histology

Tissue from the same cohort of mice used for RNAseq was also processed for histological analysis. Samples were dissected free and fixed by immersion in Bouin’s fixative overnight. Tissue was then processed by standard protocols for paraffin embedding and microtome sectioning at 4-6 microns of sections. After sectioning, tissue was mounted on slides and stained using a Leica ST5010 automated slide stainer with hematoxylin and eosin by standard protocols.

Purkinje cell density was quantified by defining a line of known length along the granule cell/molecular layer boundary in H&E stained sections and counting the number of Purkinje cell bodies per unit length. Photoreceptor cell density was quantified by defining a region of interest in the outer nuclear layer and counting the number of photoreceptor nuclei per square micron. The same analysis provided outer nuclear layer thickness, as areas were defined from the outer plexiform layer to the outer boundary of the outer nuclear layer. Retinal ganglion cell number was determined by counting the number of cells in the ganglion cell layer across a known distance in the central retina near the nerve head.

Femoral nerves were fixed by immersion in 2% paraformaldehyde and 2% glutaraldehyde in a 0.1M cacodylate buffer. Sections were then dehydrated and plastic embedded as described, and 0.5 micron sections were stained by toluidine blue (38). Images were captured with a 40X objective lens and myelinated axons were counted.

#### Neuromuscular junction staining

The plantaris muscle was fixed in freshly prepared 4% paraformaldehyde in PBS for ~4 hours at 4°C and rinsed in PBS. The tissue was then teased and stained as a wholemount for neurofilament light chain (2H3, DHSB) and SV2 (DHSB) and Alexa594 conjugated alpha-bungarotoxin as described (39).

### Data Filtering

Differentially expressed genes and TMM-normalized reads were output from a standard differential expression analysis script based on edgeR (27) implemented by the Computational Sciences Core at The Jackson Laboratory. Lists of differentially expressed genes from edgeR were filtered to remove those with false discovery rate (FDR) > 0.05. Only differentially expressed genes with an absolute log2 fold-change (logFC) ≥ 1.5 were considered in downstream analysis. Unless otherwise stated “metabolites” refer to those metabolites identified by the GCTOF platform from powdered tissue (indicated by PT) samples. Metabolites output from MetaboAnalyst5.0 comparisons were filtered to remove those with FDR > 0.05, and all changes in abundance were considered regardless of fold change value. Differentially expressed genes were compared between tissues and timepoints using the Venny2.0 or Deep Venn interactive Venn Diagram web applications (40,41).

### Annotation Enrichment Analysis

Lists of filtered differentially expressed genes were uploaded to MouseMine “list” query (42). Gene Ontology terms (biological process, cellular compartment, molecular function)(43-45), Reactome pathway ontologies (46), and Mammalian Phenotype Ontology (47) terms were ranked by p-value and exported as comma separated value (csv) files. The number of genes annotated in each database at the time of analysis (December 1, 2021) are provided in (Supp. Table 1). Significance values were -log_10_ transformed for visualization via bar plots and dot plots in R studio.

### Gene Set Enrichment Analysis

Trimmed mean of M-values (TMM) normalized counts were input to Gene Set Enrichment Analysis (GSEA) software (48). We performed comparisons against numerous gene sets from the online database, and Reactome pathways were included in the final report (46). 1500 phenotype permutations with No Collapse were entered as required parameters. Default basic fields were used with a weighted enrichment statistic and Signal2Noise used for ranking genes.

We performed a leading-edge analysis in GSEA to determine which subsets of genes contribute most to enrichment of a given process, and to find the most significant core processes by adjusted p-values, which we cross-referenced with differential expression data to improve our biological interpretation.

### NetworkAnalyst3.0

Lists of filtered differentially expressed genes from tissues where no metabolites passed FDR filtering were uploaded to NetworkAnalyst3.0 (49). Generic protein-protein interaction (PPI) networks, Transcription factor (TF)-Gene interactions, and Interactive cord diagram functionalities were used to understand gene set interaction, regulation, and similarity.

### MetaboAnalyst5.0

Metabolites that passed FDR filtering were uploaded to the Enrichment module of MetaboAnalyst5.0. The uploaded list was cross-referenced against the Metaboanalyst5.0 internally curated “Disease signatures” metabolite sets for Blood and CSF. Filtered differentially expressed genes and metabolites lists were uploaded to MetaboAnalyst5.0 Joint-Pathway module using the “Metabolic pathways” setting and default parameters (50,51). A comprehensive evaluation of MetaboAnalyst5.0 parameterization on our dataset revealed no influence of parameter variation on the top identified pathways (those pathways meeting FDR and Adj. p-value cutoff ≤ 0.05 cutoff). Pathways that passed FDR and Adj. p-value cutoffs were ranked by impact score. Lists of metabolites and differentially expressed genes implicated in top pathways were then projected onto KEGG pathway maps using MetaboAnalyst5.0 internal function.

### Data Visualization in GraphPad Prism and R Studio

Ranked ontologies from NetworkAnalyst3.0, MetaboAnalyst5.0, and MouseMine were visualized as bar plots using GraphPad Prism 8 or as dot plots made with GGPlot2 for R statistical software in R Studio (52,53). TMM normalized counts were visualized using the Pheatmap package for R studio (54). Clusters were identified in a supervised manner using dendrogram height (h=15, k=4) to separate genes into four categories based on expression level and consistency of expression between samples. Ontologies for each clustered were queried in MouseMine as above, and grouped into major categories: neuronal function, nervous system structure, behavior, seizures, movement, and other. Category membership is visualized using circular plots. Gene-disease associations are visualized using the enrichplot R package (55). NCBI gene IDs were mapped to Ensembl IDs (GRCm38.78) from differential expression analysis pipeline using Ensembl Biomart.

### Targeted Pathway Mapping

Lists of differentially expressed genes identified by Ensembl IDs (GRCm38.78) were filtered to remove those with FDR > 0.05, and NCBI GeneIDs were assigned to each using the KEGG Mapper – Convert ID function. NCBI GeneID lists were then entered into the KEGG Mapper – Search Pathway module using the organism-specific search mode (set to *Mus musculus*) (56). Mappings of our gene lists to Lysine Degradation and Fatty Acid Degradation pathways were exported.

## Acknowledgements and Funding

The authors would like to thank the Scientific Services at The Jackson Laboratory for their technical assistance in this project, including the Fine Mapping service, the Histology and Electron Microscopy service, the Genome Technologies service, and the Computational Sciences service. The services are a Shared Resource of the JAX Cancer Center (P30 CA034196). This work was supported by the National Institutes of Health: R21 NS082666, R37 NS054154 to RWB, and R01 EY011721 to SWMJ. SWMJ was also supported by funds from the RBP, Columbia University. The West Coast Metabolomics Center is supported by U2 ES030158. GCM was supported by T32 GM132006.

---

**Supplemental Figure 1.**
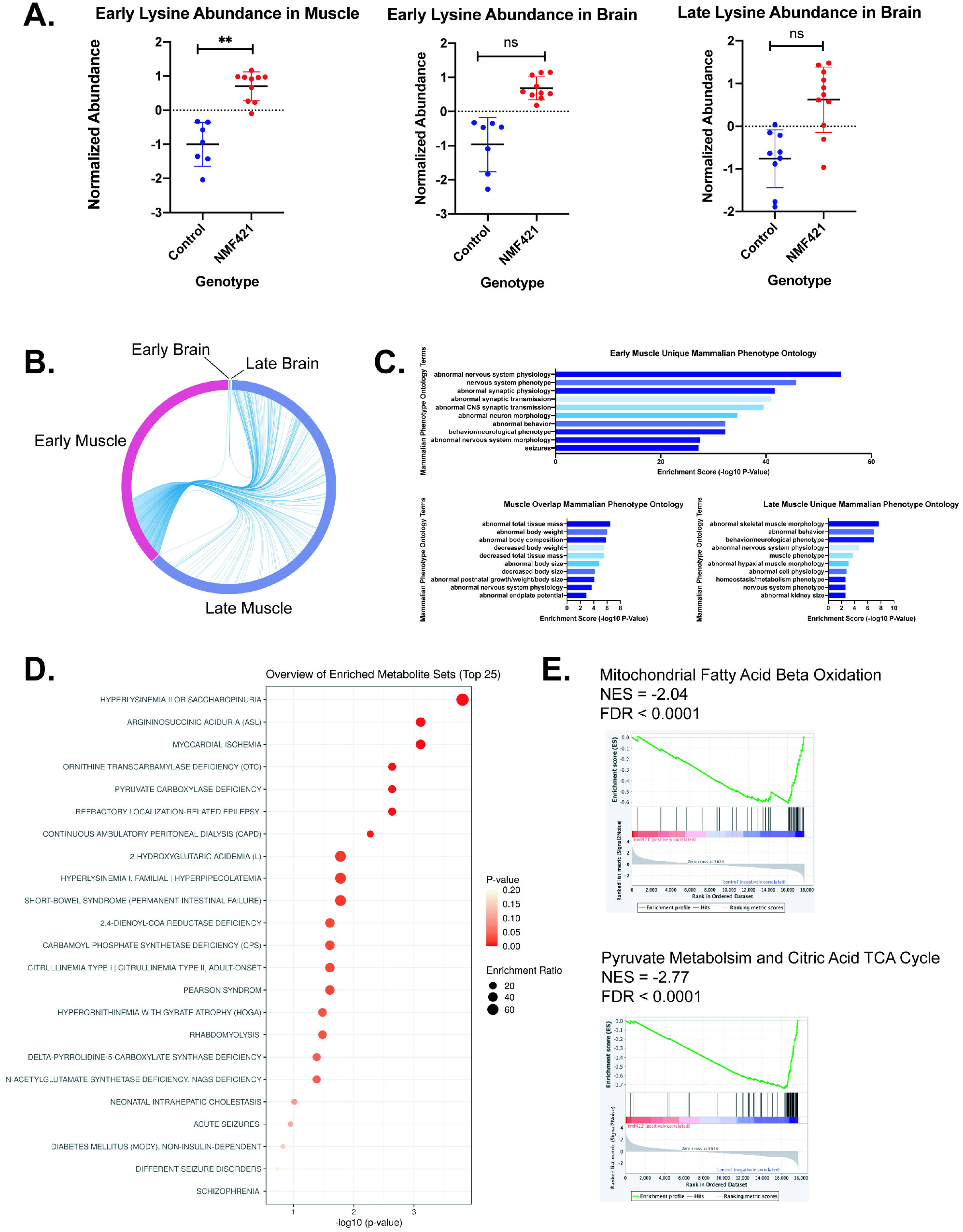
Further transcriptomic and metabolomic analysis of *Nadk2* mutant tissue. A) Abundance of lysine in early timepoint muscle and brain at early and late timepoints of *Nadk2^-/-^* mice. Significance is determined by t-test in MetaboAnalyst5.0. B) Cord diagram depicting overlapping differentially expressed genes in *Nadk2^-/-^* muscle and brain at early and late timepoints. C) Mammalian Phenotype Ontology terms associated with differentially expressed genes that are specific to muscle at the early timepoint, shared between muscle at the early and late timepoints, and specific to muscle at the late timepoint. D) Human conditions associated with the list of metabolites with significant differences in abundance between *Nadk2^-/-^* muscle and wildtype mice at the early timepoint in MetaboAnalyst5.0. Color indicates p-value and dot size represents the enrichment ratio. E) Gene set enrichment analysis (GSEA) diagrams for key *Nadk2* deficiency pathways in control relative to *Nadk2^-/-^* muscle at the late timepoint.

**Supplemental Figure 2.**
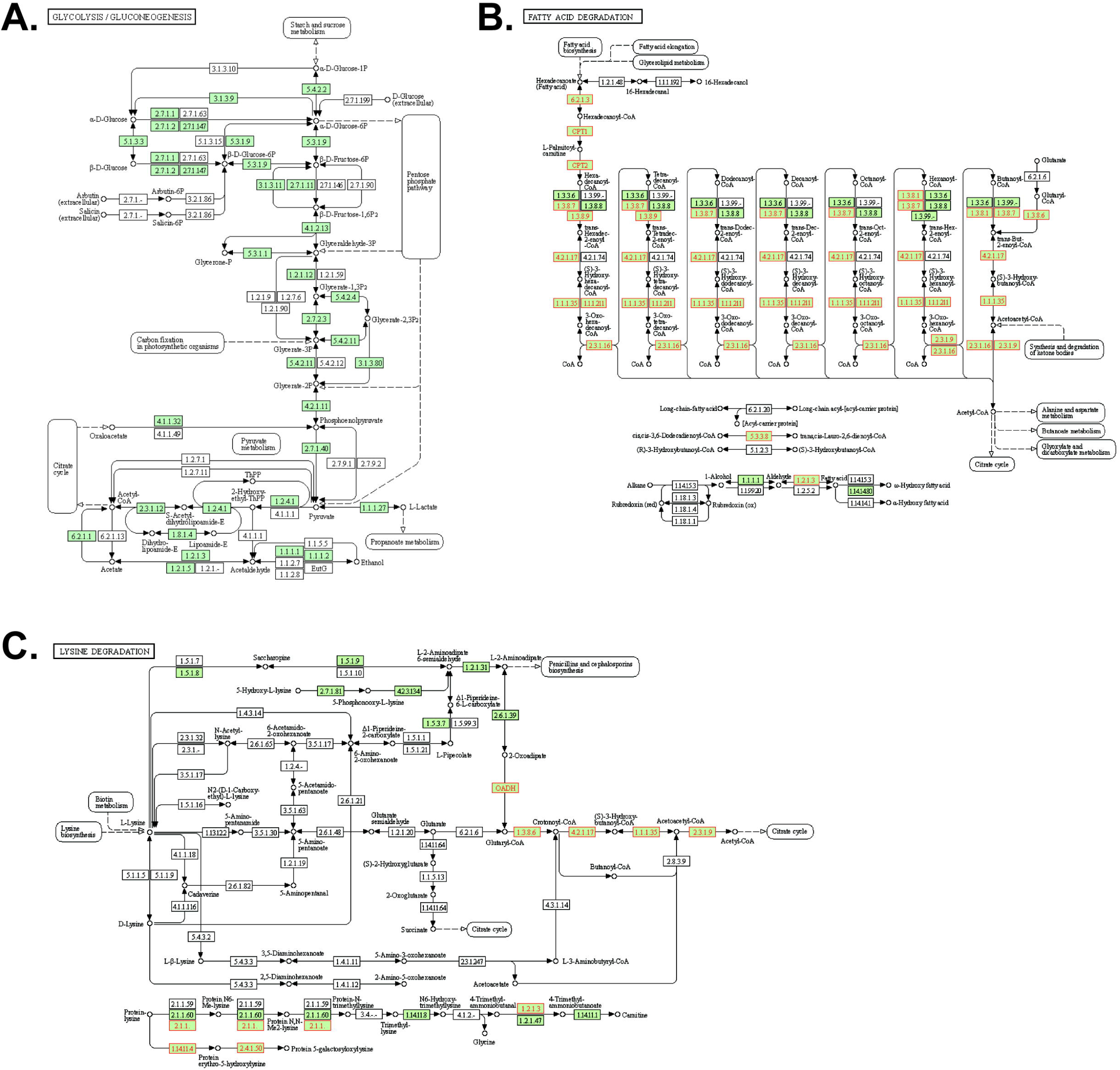
KEGG map overlaid with differentially expressed genes from *Nadk2^-/-^* late muscle tissue. A) Differentially expressed genes and metabolites uploaded to MetaboAnalyst5.0 are overlaid on KEGG glycolysis and gluconeogenesis pathway. Green indicates pathway components represented in the mouse genome. B) The KEGG mapper function was used to overlay differentially expressed genes from *Nadk2^-/-^* late muscle tissue on the canonical lysine degradation pathway. Red indicates differentially expressed genes that overlapped with pathway constituents. C) The KEGG mapper function was used to overlay differentially expressed genes from *Nadk2^-/-^* late muscle tissue on the canonical lysine degradation pathway. Red indicates differentially expressed genes that overlapped with pathway constituents.

**Supplemental Figure 3.**
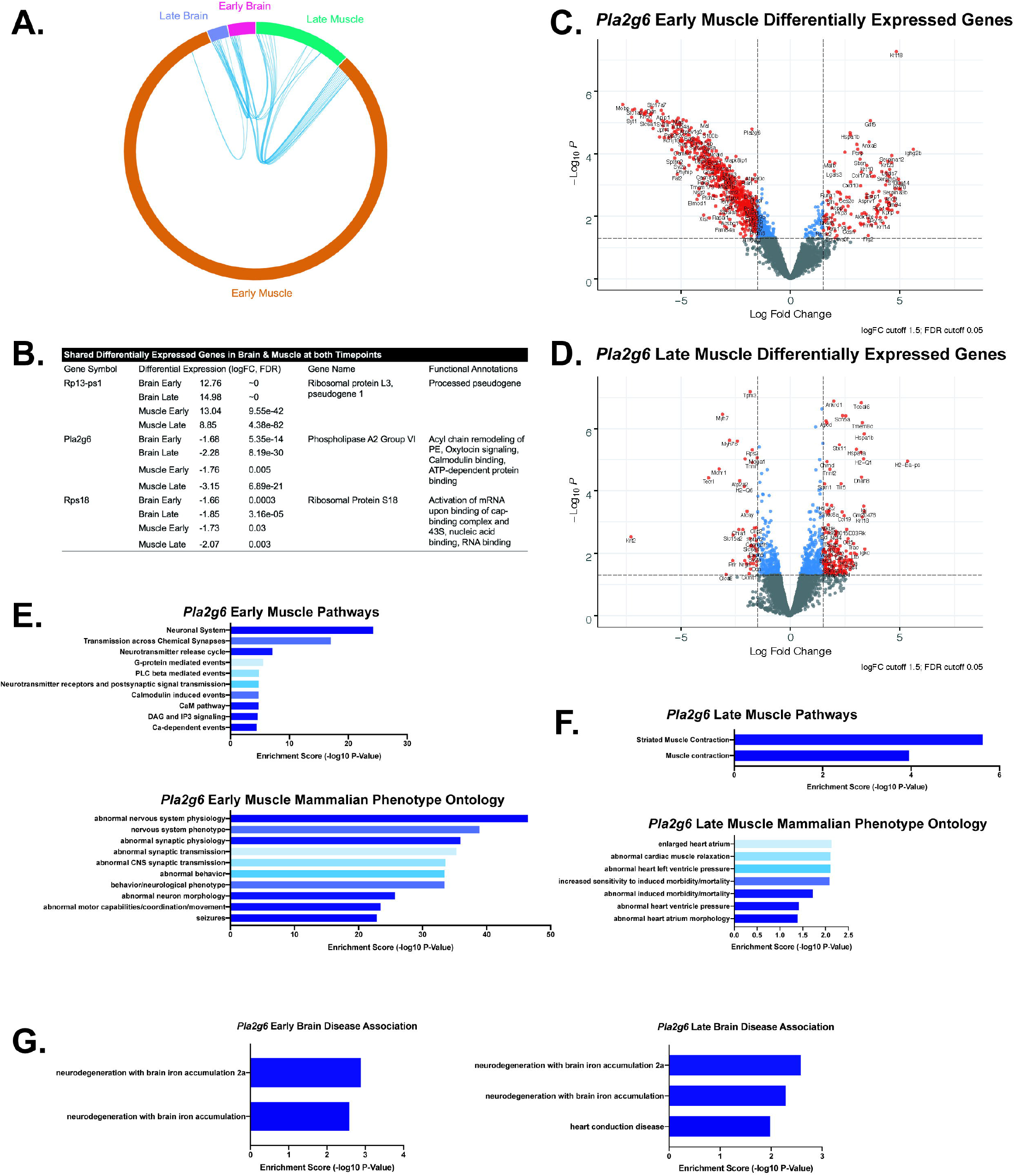
Transcriptomic analysis of *Pla2g6* muscle tissue. A) Cord diagram depicting differentially expressed genes from brain and muscle at early and late timepoints of *Pla2g6^-/-^* mice. B) Table with names, differential expression information, and functional annotations of the three differentially expressed genes that are shared between brain and muscle at both timepoints. C) Volcano plot of differentially expressed genes in *Pla2g6^-/-^* muscle at the early timepoint. Genes charted in red were considered in our analysis. D) Volcano plot of differentially expressed genes in *Pla2g6^-/-^* muscle at the late timepoint. Genes charted in red were considered in our analysis. E) Pathways and Mammalian Phenotype Ontology terms associated with differentially expressed genes from *Pla2g6^-/-^* muscle at the early timepoint. F) Pathways and Mammalian Phenotype Ontology terms associated with differentially expressed genes from *Pla2g6^-/-^* muscle at the late timepoint. G) Human diseases associated with differentially expressed genes in *Pla2g6^-/-^* brain at early and late timepoints. Note, for this set nominal p-values are used. Significance threshold was not reached after Holm-Bonferroni post-hoc test.

